# German *Ixodes inopinatus* samples may not actually represent this tick species

**DOI:** 10.1101/2023.02.14.528458

**Authors:** Robert E. Rollins, Gabriele Margos, Andreas Brachmann, Stefan Krebs, Alexia Mouchet, Niels J. Dingemanse, AbdElkarim Laatamna, Nassiba Reghaissia, Volker Fingerle, Dirk Metzler, Noémie S. Becker, Lidia Chitimia-Dobler

**Author notes:** Corresponding Author: Robert E. Rollins, Institute of Avian Research “Vogelwarte Helgoland”, An der Vogelwarte 21, 26386 Wilhelmshaven, Germany. These authors share senior authorship for this manuscript (Noémie S. Becker, Lidia Chitimia-Dobler).

## Abstract

Ticks are important vectors of human and animal pathogens, but many questions remain unanswered regarding their taxonomy. Molecular sequencing methods have allowed research to start understanding the evolutionary history of even closely related tick species. *Ixodes inopinatus* is considered a sister species and highly similar to *Ixodes ricinus*, an important vector of many tick-borne pathogens in Europe, but identification between these species remains ambiguous with disagreement on the geographic extent of *I. inopinatus*. In 2018-2019, 1583 ticks were collected from breeding great tits (*Parus major*) in southern Germany, of which 45 were later morphologically identified as *I. inopinatus*. We aimed to confirm morphological identification using molecular tools. Utilizing two genetic markers (16S rRNA, TROSPA) (n=37) and whole genome sequencing of specific ticks (n=8), we were able to determine that German samples morphologically identified as *I. inopinatus*, genetically represent *I. ricinus* regardless of previous morphological identification and most likely are not *I. ricinus*/*I. inopinatus* hybrids. Further, our results showed that the entire mitochondrial genome, let alone singular mitochondrial genes (i.e., 16S), is unable to distinguish between *I. ricinus* and *I. inopinatus*. As most examples of *I. inopinatus* in Germany were based on morphology and mitochondrial sequences, the results of the current study brings into question whether *I. inopinatus* was properly identified in previous research and if this species exists in Central Europe. Our results highlight the power of utilizing genomic data in answering questions regarding tick taxonomy even when closely related species are considered.

**Highlights:**

- German *Ixodes inopinatus* samples represent *I. ricinus* based on genomic data
- The mitochondrial genome is not sufficient for delineation of *I. inopinatus* and *I. ricinus*
- German samples most likely do not represent *I. ricinus*/*I. inopinatus* hybrids

## 1 Introduction

Ticks are parasitic arthropods belonging to the order Acari along with other species of mites and can be found on all continents besides Antarctica (Estrada-Peña et al., 2004; Hillyard, 1996). Additionally, ticks act as major vectors of pathogens to domestic animals and humans worldwide, rivaled only by mosquitos in their medical importance (Hillyard, 1996). Due to their importance in human and animal disease, it is of utmost importance to have clear and unambiguous methods for identification of tick species even when closely related species are under study. Even so, many species remain challenging to identify without expert opinion with even this resulting in ambiguous identification of specimens in some cases. This highlights the need for methods which identify tick specimens unambiguously so as to aid in the study of their evolutionary history.

Many tick species have been and still are identified based on morphological characteristics (Estrada-Peña et al., 2004; Guglielmone et al., 2014; Hillyard, 1996). Progress in molecular sequencing tools and methods/technologies, though, has increased the number of studies using different genetic markers for confirmation of morphological identification (Abouelhassan et al., 2019) and also further describing the evolutionary history of various tick species (Charrier et al., 2019; Wang et al., 2019; Xu et al., 2003). Many genetic markers used for tick species identification are located on the mitochondrial genome, which has been widely and preferentially used, due to its overall conservation, ease of amplification, and existing as a haploid sequence (Abouelhassan et al., 2019). On another side, more recent studies have started to also utilize genetic data from the nuclear genome, results of which highlight differences that were not apparent based on mitochondrial studies alone (Charrier et al., 2019; Jia et al., 2020; Poli et al., 2020). Even so, the tick genome has proved complex for genome assembly due to its large size (>2Gb) and high variability in chromosome structure, resulting in very few published reference genomes (Cramaro et al., 2017; Gulia-Nuss et al., 2016; Jia et al., 2020). Even so, nuclear based genomic studies have showed the ability of these methods to unravel the evolutionary history of even closely related tick species (Jia et al., 2020; Poli et al., 2020), opening up the possibility to study the taxonomy and identification of other closely related species such as in the case of *Ixodes ricinus* Linnaeus, 1758 and *Ixodes inopinatus* Estrada-Peña, Petney, Nava, 2014.

*Ixodes ricinus* is one of the main vectors of various tick-borne pathogens such as tick-borne encephalitis virus and *Borrelia burgdorferi* sensu lato (Hillyard, 1996). Based on genetic data, it has been shown that this tick species forms two distinct populations between Europe and North Africa (Noureddine et al., 2011; Poli et al., 2020). Subsequent research supported that these distinct clusters represented two separate tick species, *Ixodes inopinatus* in North Africa and *I. ricinus* in Europe (Estrada-Peña et al., 2014; Poli et al., 2020). It is now believed that both species exist in sympatry across their range (Chitimia-Dobler et al., 2018; Rubel et al., 2021; Younsi et al., 2020). Ambiguity in morphological characteristics between *I. ricinus* and *I. inopinatus* requiring expert knowledge to delineate (Estrada-Peña et al., 2014) and recent challenges to use certain common genetic markers for species determination (Plantard et al., 2022) have complicated the unambiguous identification and study of *I. inopinatus*. This fact paired with recent work reporting *I. inopinatus* samples positive for tick-borne pathogens (i.e., *B. burgdorferi* sensu lato) (Hauck et al., 2020; Knoll et al., 2021) and movement on migratory birds (Toma et al., 2021) further underlies the need for a clear methodology to identify *I. inopinatus* so as to facilitate research into the overall biology of this tick species.

In the years 2018-2019, several (n=45) larvae and nymphs were collected as part of a previous project from breeding great tits (*Parus major*) in various nest-box plots located south of Munich, Germany and morphologically identified as *I. inopinatus* (Rollins et al., 2021). As *I. inopinatus* had previously not been observed on this bird species, we aimed to first determine if the samples that were morphologically identified as *I. inopinatus* truly represented this species. Using genetic markers (16S rRNA, TROSPA) on all tick specimens (n=37) and whole genome sequencing of specific samples (n=8), we are able to show here that samples morphologically identified as *I. inopinatus* most likely represent *I. ricinus* regardless of morphological identification. These results, paired with phylogenetic and demographic estimation based on whole genome sequencing, suggests that samples identified as *I. inopinatus* previously in Germany (Chitimia-Dobler et al., 2018; Hauck et al., 2020; Knoll et al., 2021; Rollins et al., 2021) may not actually represent this tick species and that *I. inopinatus* may be limited to Northern Africa and, potentially, southern Spain.

## 2 Methods

### 2.1 Samples, DNA extraction, and PCR analysis

For this study, 45 samples morphologically identified as *I. inopinatus* (31 nymphs and 14 larvae) were used from a previous study (Rollins et al., 2021). These ticks were each collected feeding on breeding great tit adults (*Parus major*) south of Munich, Germany in the years 2018-2019. Additionally, three German nymphs from the same project morphologically identified as *I. ricinus*, and two adult, male ticks collected from cattle in the Ain Sandel district (Guelma province) of eastern Algeria and morphologically identified as *I. inopinatus* were included as controls. All ticks were stored in 99% ethanol prior to morphological identification and molecular analyses. Morphological identification was performed according to published taxonomic keys (Estrada-Peña et al., 2014; Filippova, 1977; Hillyard, 1996).

We first aimed to confirm morphological identification performed in Rollins et al., (2021), for which we chose to amplify one mitochondrial gene (16S rRNA) and one nuclear gene (TROSPA) through PCR as these markers have been shown to separate *I. ricinus* and *I. inopinatus* in previous work (Estrada-Peña et al., 2014; Noureddine et al., 2011). Tick DNA was extracted either using the BioSprint All Wet Vet Kit (Qiagen, Hilden, Germany) or a Maxwell® 16 LED DNA kit (Promega, Madison, WI, USA) according to standard procedure. Extracted DNA was stored at -20°C prior to further analysis. To amplify each gene, published primers and protocols were used as described by Mangold et al., (1998) for 16S rRNA and Noureddine et al. (2011) for TROSPA. Individual PCR products were sequenced on an ABI3730 (Applied Biosystems, USA) at the Genomics Service Unit (LMU Biocenter, Munich, Germany). Quality of sequencing runs were checked manually in FinchTV Version 1.4.0 (Geospiza, Inc.; Seattle, WA, USA; http://www.geospiza.com). Heterozygote positions were allowed for all TROSPA sequences and we encoded as the IUPAC ambiguity codes. For 16S rRNA sequences, ambiguous base calls were encoded with N as this sequence is haploid and therefore cannot have heterozygote positions. Low quality bases (called as N) at the beginning and end of sequences were removed prior to further analysis.

### 2.2 Median-joining networks

Sequences for each gene were compiled into files containing reference sequences for *I. ricinus* and *I. inopinatus* based on previous research (16S rRNA, GenBank PopSet: 309318023; TROSPA, GenBank PopSet; 309318631 and 309318389) (Noureddine et al., 2011; Poli et al., 2020). These sequences were then aligned using MUSCLE v3.8.425 (Edgar, 2004a, 2004b) as implemented in Aliview v1.28 (Larsson, 2014). Clustering based on both 16S rRNA and TROSPA genes was determined through reconstructing median-joining networks (MJN) calculated in Network v5.0.1.1. (Fluxus Technology Ltd., Stanway, England). Haplotype RTF files were generated by DnaSP v.6 without considering gaps or missing data and removed invariable sites (Rozas et al., 2017). Only for the case of TROSPA, sequences were first phased using PHASE v.2.1 (Stephens et al., 2001; Stephens and Donnelly, 2003) as implemented in DnaSP v.6 (Rozas et al., 2017) prior to haplotype determination and MJN analysis.

### 2.3 Whole genome sequencing

In the clustering analysis, we observed discrepancies between morphological and molecular species identification, especially in the case of German samples that were morphologically identified as *I. inopinatus*. To definitively determine if the German samples represented *I. inopinatus*, we chose to produce whole genome sequencing data for eight individual ticks (see Table 1). Ticks were chosen based on the three criteria used to sort samples to *I. inopinatus* or *I. ricinus* in this study: morphology, 16S rRNA sequence, and TROSPA sequence. *Ixodes inopinatus* samples were chosen based on meeting one, two, or all three criteria with two *I. ricinus* (morphology, 16S rRNA, and TROSPA) being included as well. For full information on each sample see Table 1.

**Table 1.** Meta-data of tick samples used for whole genome sequencing.

| ID | Year | Country | Source | Life-Stage | Morphology | 16S rRNA | TROSPA |
| --- | --- | --- | --- | --- | --- | --- | --- |
| **ALG1** | 2021 | Algeria | Cattle | Adult/Male | *Ixodes inopinatus* | *Ixodes inopinatus* | *Ixodes inopinatus* |
| **ALG2** | 2021 | Algeria | Cattle | Adult/Male | *Ixodes inopinatus* | *Ixodes inopinatus* | *Ixodes inopinatus* |
| **8-C12** | 2018 | Germany | Parus major | Nymph | *Ixodes inopinatus* | *Ixodes inopinatus* | *Ixodes ricinus* |
| **11-E12** | 2018 | Germany | Parus major | Nymph | *Ixodes inopinatus* | *Ixodes inopinatus* | *Ixodes ricinus* |
| **6-F6** | 2018 | Germany | Parus major | Nymph | *Ixodes inopinatus* | *Ixodes ricinus* | *Ixodes ricinus* |
| **12-E9** | 2018 | Germany | Parus major | Nymph | *Ixodes inopinatus* | *Ixodes ricinus* | *Ixodes ricinus* |
| **12-F9** | 2018 | Germany | Parus major | Nymph | *Ixodes ricinus* | *Ixodes ricinus* | *Ixodes ricinus* |
| **3-F8** | 2018 | Germany | Parus major | Nymph | *Ixodes ricinus* | *Ixodes ricinus* | *Ixodes ricinus* |

For extracted DNA used in PCR analysis, quality (260/280) and concentration were measured using a NanoDrop^®^ 1000 photometer (Thermo Fisher Scientific, Waltham, MA, USA) and a Qubit^®^ 3.0 fluorometer (Thermo Fisher Scientific, MA, USA), respectively. For all German tick samples (*I. inopinatus, I. ricinus*) libraries were produced using the 1SPlus library preparation kit (Swift Biosciences, Ann Arbor, USA) with DNA fragmented to an approximate size of 300 bp with a Covaris M220 sonicator (Covaris,Woburn, USA). Libraries were dual-indexed, pooled and sequenced on a NovaSeq 6000 sequencer (Illumina, San Diego, CA, USA; 2×150 bp paired-end sequencing) at Novagene Europe (Cambridge, UK). For both Algerian *I. inopinatus* samples, libraries were produced from 50 ng of DNA with the Nextera XT DNA Sample Preparation Kit (Illumina, Germany) according to the manufacturer’s protocol. The library was quality controlled by analysis on an Agilent 2000 Bioanalyzer with the Agilent High Sensitivity DNA Kit (Agilent Technologies, Santa Clara, CA, USA) for fragment sizes of ca. 300-600 bp. Sequencing on a MiSeq sequencer (Illumina, San Diego, CA, USA; 2×250 bp paired-end sequencing, v3 chemistry) was performed in the Genomics Service Unit (LMU Biocenter, Martinsried, Germany). Additional sequencing was performed on a NovaSeq 6000 sequencer (Illumina, San Diego, CA, USA; 2×150 bp paired-end sequencing) at Novagene Europe (Cambridge, UK).

### 2.4 Assembly of mitochondrial genomes and nuclear gene sequences

Prior to analysis, Illumina reads were first trimmed for Illumina NovaSeq or MiSeq adapter sequences dependent on sequencing technology using Trimmomatic v. 0.38 (Bolger et al., 2014a, 2014b). To produce full mitochondrial genomes, raw reads were mapped to the *I. ricinus* mitochondrial reference genome (GenBank: NC_018369.2) using BWA-MEM v0.7.17 (Li, 2013; Li and Durbin, 2009). Mapping results were filtered based on the following criteria: mapping quality >= 20, minor allele frequency (MAF) >= 0.2, and a minimum of 10 counts at each site. After mapping, whole mitochondrial genome sequences were reconstructed for each sequenced tick based on the mapping results: for each variable position the alternative SNP was called if the alternative frequency was greater than or equal to 70% whereas the reference SNP was called if the alternative frequency was less than or equal to 30%, otherwise the position was called as N.

Nuclear gene sequences were reconstructed in a similar manner by mapping to the *I. ricinus* reference genome (GenBank: GCA_000973045.2), but with a few key differences. Raw reads were mapped using BWA-MEM v0.7.17 (Li, 2013; Li and Durbin, 2009) to the reference sequences. From these results, a random 1000 bp long sequence was chosen from each of the scaffolds from the reference that were over 10 kb in length. Sequences were only kept if at least 950 base pairs of the sequence had at least a coverage of 10x. Sequences were then reconstructed based on the following rules: homozygote for alternative allele if over 80% of reads bear alternative allele (minimum of five reads), heterozygote if 20-80% of reads bear alternative allele (minimum of five reads), otherwise homozygote for reference allele. In total, 435 sequences were identified through this mapping analysis. To obtain an outgroup for demographic and phylogenetic analysis, these 435 sequences were then compared to the *I. scapularis* reference genome (GenBank: GCA_016920785.2) using BLAST v.2.8.1 (Altschul et al., 1990; Camacho et al., 2009) (algorithm: *blastn*). Blast hits were kept only if they showed a single hit which displayed 80% identify over 80% of the entire sequence. From this analysis, 194 sequences were identified to be used in further analyses.

### 2.5 Phylogenetic analyses

The mitochondrial genome has been shown recently to be more powerful in terms of species delineation than single gene markers (Wang et al., 2019). For this reason, we reconstructed the phylogenetic tree of our tick samples using the full mitochondrial genome sequence. Mitochondrial genome sequences for our eight sequenced tick specimens were aligned with the *I. ricinus* reference (GenBank: NC_018369.2) using MUSCLE v3.8.425 (Edgar, 2004a, 2004b) as implemented in Aliview v1.28 (Larsson, 2014). The *Ixodes persulcatus* mitochondrial genome (GenBank: NC_004370.1) was additionally included as an outgroup. The phylogeny was then estimated in BEAST v2.6.6 (Bouckaert et al., 2019) with the following parameters: gamma site model with four categories, log-normal relaxed clock (Drummond et al., 2006), GTR substitution model (Tavaré, 1986). Three independent runs were launched for 50 million generations each. Convergence was checked with Tracer v. 1.7.1 (Rambaut et al., 2018). The best tree was selected using TreeAnnotator v. 1.10.4 (Drummond and Rambaut, 2007) with a relative burn-in of 10% for two runs and a burn-in of 50% for the third. Convergence to a single topology in all three independent runs was checked manually in FigTree v. 1.4.4 (http://tree.bio.ed.ac.uk/software/figtree/), which was also used to plot the tree shown in Figure 2A.

We additionally aimed to reconstruct the phylogeny of our samples based on the 194 nuclear sequences extracted from the whole genome sequencing data produced during this study. For this, we analyzed unphased sequences (n=194) for all eight sequenced ticks and included sequences identified in both the *I. ricinus* (GenBank: ASM97304v2) and *I. scapularis* (GenBank: GCA_016920785.2) reference genomes which were individually aligned using MUSCLE v3.8.425 (Edgar, 2004a, 2004b). Phylogeny reconstruction was then performed in MrBayes v3.2.6 as a partitioned analysis (Huelsenbeck and Ronquist, 2001; Ronquist et al., 2012). For this, each sequence was allowed to evolve independently by estimating a GTR (Tavaré, 1986) substitution model with inverse gamma distributed rate variation and ploidy set to diploid per individual gene. Three independent runs were launched for 100 million generations at which point convergence was checked with Tracer v. 1.7.1 (Rambaut et al., 2018). Consensus trees were produced by the *sumt* command in MrBayes with a relative burn-in of 25%. Convergence to a single topology in all three independent runs was checked manually in FigTree v. 1.4.4 (http://tree.bio.ed.ac.uk/software/figtree/), which was also used to plot the tree shown in Figure 2B.

### 2.6 Estimating heterozygosity and demographic history

Heterozygosity for all sequenced tick samples was estimated in R v.3.5.2 (R Core Team, 2019) using the package GENHET (Coulon, 2010).

Demographic estimates were produced using jaatha 3.2.3, https://CRAN.R-project.org/package=jaatha (Mathew et al., 2013; Naduvilezhath et al., 2011) also using the 194 unphased nuclear sequences which included an outgroup (i.e., *I. scapularis*). The model assumed three populations that all shared a common ancestor. Population 1 contained the German *I. ricinus* samples (12-F9, 3-F8), Population 2 contained all morphologically identified *I. inopinatus* samples from Germany (8-C12, 11-E12, 6-F6, 12-E9), and Population 3 contained the Algerian *I. inopinatus* samples (ALG1, ALG2). We allowed for gene flow between populations 2 and 3 (those morphologically identified as *I. inopinatus*) as well as between populations 1 and 2 as these two German populations could represent a larger central-European population. We did not allow direct gene flow between populations 1 and 3. The mutation rates between populations *i* and *j* are given as *M*_*ij*_=4*N*_2_*m*_*ij*_, where *N*_2_ is the effective size of population 2 and *m*_*ij*_ is the fraction of individuals in population *i* that are immigrants from population *j*. We assumed that all sampled loci were diploid, such that an effective population size of *N*_2_ individuals corresponds to an effective population size of 2*N*_2_ alleles. As for gene flow, we did not allow the possibility for population split events between populations 1 and 3 neglecting that these populations could be more closely related to each other than to population 2. Therefore, we assumed that either the two European populations (Pop 1, Pop 2) stem from an ancestral population from which the Northern African population (Pop 3) has branched off further back in time or that the two *I. inopinatus* populations (Pop 2, Pop 3) recently split from a joint ancestral population and that this joint ancestral population has a joint ancestral population with the *I. ricinus* population (Pop 1). The time span since the split between populations 1 and 2 (or the joint ancestor of 2 and 3) is *t*_12_ and the time since the split between populations 2 and 3 (or the ancestral population of 1 and 2) is *t*_23_, where time is measured in units of 4*N*_2_ generations. For the ancestral populations we assumed the same population size as for population 2. Besides the migration rates and the population split times, further model parameters are the population-migration rate θ=4*N*_2_*μ* of population 2 where *μ* is the mutation rate per generation and locus (of 1000 bp), and the size ratios between populations 1 and 2 and between populations 2 and 3. Finally, ρ is the recombination rate per locus and per 4*N*_2_ generations and, as we assumed an HKY finite-sites mutation model, we estimated the transition–transversion ratio of mutations. See Figure S1 for a schematic overview of the demographic model. We chose the populations with the *a priori* expectation that morphological identification is correct, and set up the model to determine if the demographic estimates support or refute this assumption. Additionally, by including the assumed German *I. inopinatus* samples as their own population and allowing for gene flow with German *I. ricinus* and Algerian *I. inopinatus* we can assess gene flow between the populations to determine if the assumed German *I. inopinatus* samples potentially represent hybrids (i.e., equivalent gene flow from Pop 1 and Pop 3).

We used the R package coala, version 0.7.1, to specify and simulate the demographic model in R (Staab and Metzler, 2016) (https://CRAN.R-project.org/package=coala) combined with the *scrm* package (version 1.7.4) of the seq-gen program, version 1.3.4 to generate sequences evolving along the simulated genealogies according to the HKY sequence-evolution model (Rambaut and Grassly, 1997; Staab et al., 2015). Confidence intervals (CIs) were calculated from 50 parametric bootstrap repetitions (Efron and Tibshirani, 1993; Mathew et al., 2013) applying the reverse-percentile intervals method as implemented in the R command boot.ci with option type=‘basic’ (boot package version 1.3-28.1). Prior to analysis, sequence data were checked for linkage (see Supplementary Methods).

## 3 Results

### 3.1 Morphological identification of German samples does not match molecular data

High quality 16S rRNA and TROSPA sequences were both obtained for 37 out of 45 ticks under study. Samples containing one of the sequences or none were excluded from further analysis. For all *I. ricinus* and Algerian *I. inopinatus* samples included as controls high quality 16S and TROSPA sequences were obtained. Both median-joining networks supported two haplotype clusters predominantly corresponding to *I. ricinus* and *I. inopinatus* (Figure 1). However, in the 16S MJN there were GenBank references that clustered with the incorrect haplotype (Figure 1A). As previously reported, there are only two fixed nucleotide differences between the two species on the 16S rRNA sequences (Figure S2), therefore explaining that the branch separating the two haplotype clusters was shorter on the 16S rRNA gene (Figure 1A) in comparison to the nuclear TROSPA gene (Figure 1B). This would suggest more variation separating the groups along the TROSPA gene. From the German samples identified morphologically as *I. inopinatus*, only three samples clustered with GenBank references based on the 16S rRNA gene sequences (Figure 1A). None of these German samples clustered with reference *I. inopinatus* based on the TROSPA sequences (Figure 1B). Control ticks, both Algerian *I. inopinatus* and German *I. ricinus*, clustered within their respective species based on sequences of both genes (Figure 1).

**Figure 1.**
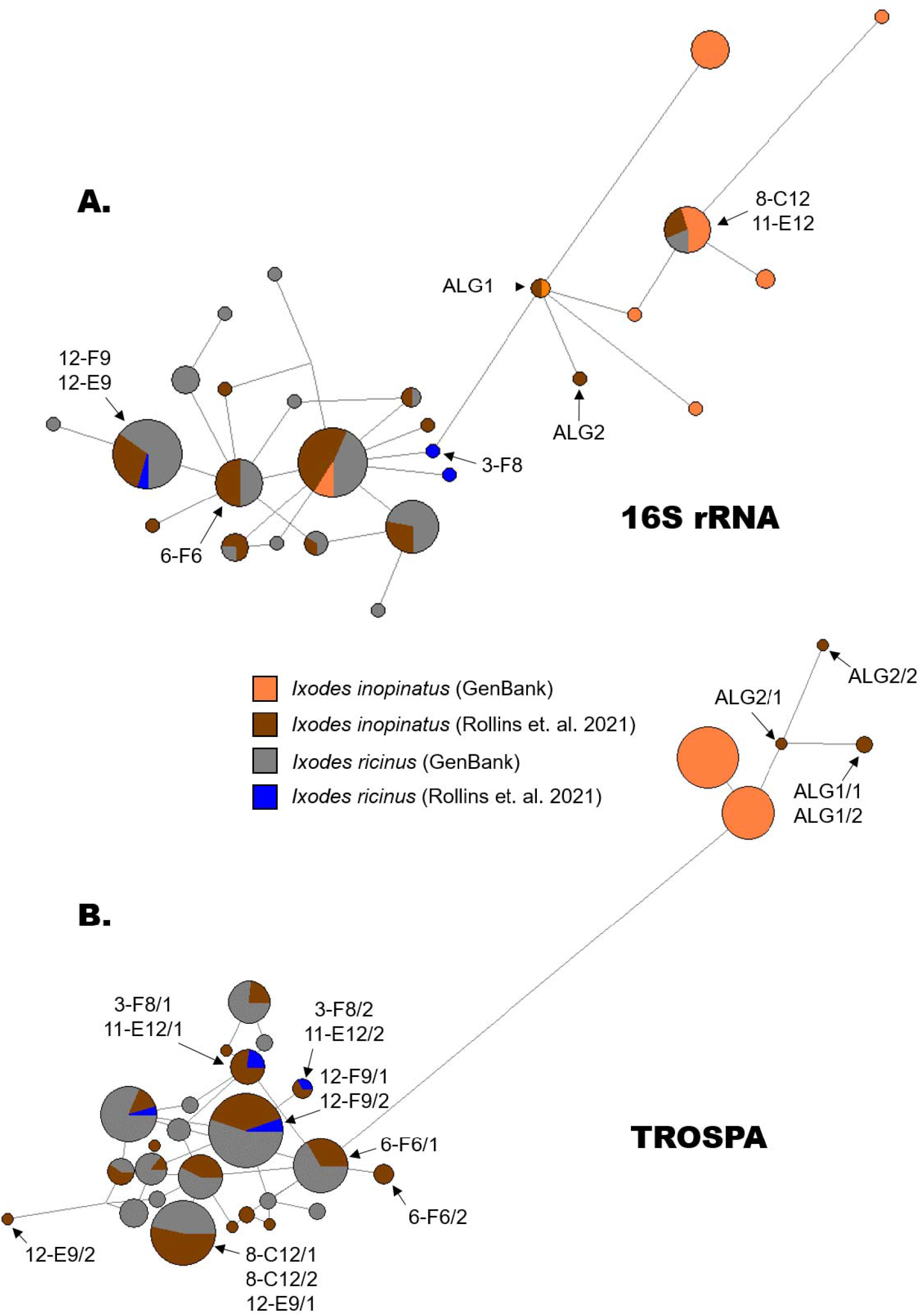
Median-joining networks (MJN) calculated in Network 5.0.1.1. (Fluxus Technology Ltd., Stanway, England) using haplotype RTF files generated by DnaSP v.6 which did not consider gaps or missing data and removed invariable sites (Rozas et al., 2017) based on either the 16S rRNA (A) or TROSPA (B) genes. For TROSPA, sequences were first phased using PHASE v.2.1 (Stephens et al., 2001; Stephens and Donnelly, 2003) as implemented in DnaSP v.6 (Rozas et al., 2017) prior to haplotype determination and MJN analysis. Arrows and sample names display the haplotype locations of samples for which we produced whole genome sequencing data.

### 3.2 Nuclear and mitochondrial phylogenies display different topologies

On average 9.04×10^8^ reads (range: 4.3×10^8^ – 1.15×10^9^ reads) accounting for an average of 135.6G (range: 65.1-172.7G) were produced per sequenced tick sample. Mitochondrial genome coverage ranged from 1890x (11-E12) to 7947x (3-F8) and no more than 31 positions displayed <10x coverage (0.21%). For all sequencing related data see Table S1. The phylogeny based on the full mitochondrial genome did not show distinct clustering between *I. inopinatus* and *I. ricinus* samples (Figure 2A). All sequenced samples (*I. inopinatus* and *I. ricinus*) formed a single clade with the *I. ricinus* reference being basal to the rest of the samples (Figure 2A). The two German samples morphologically identified as *I. inopinatus* (8-C12, 11-E12), that additionally clustered with GenBank *I. inopinatus* reference sequences of the 16S rRNA gene, as well as the Algerian *I. inopinatus* samples did not form a monophyletic clade within the larger *I. ricinus* clade (Figure 2A). The phylogeny based on 194 nuclear sequences from across the genome, however, showed two distinct clades with the Algerian *I. inopinatus* samples forming a sister clade to all other samples (Figure 2B). The *I. ricinus* reference was also nested within this second monophyletic clade of all German samples and there was no clear structuring between German samples that were originally morphologically identified as *I. inopinatus* (Figure 2B).

**Figure 2.**
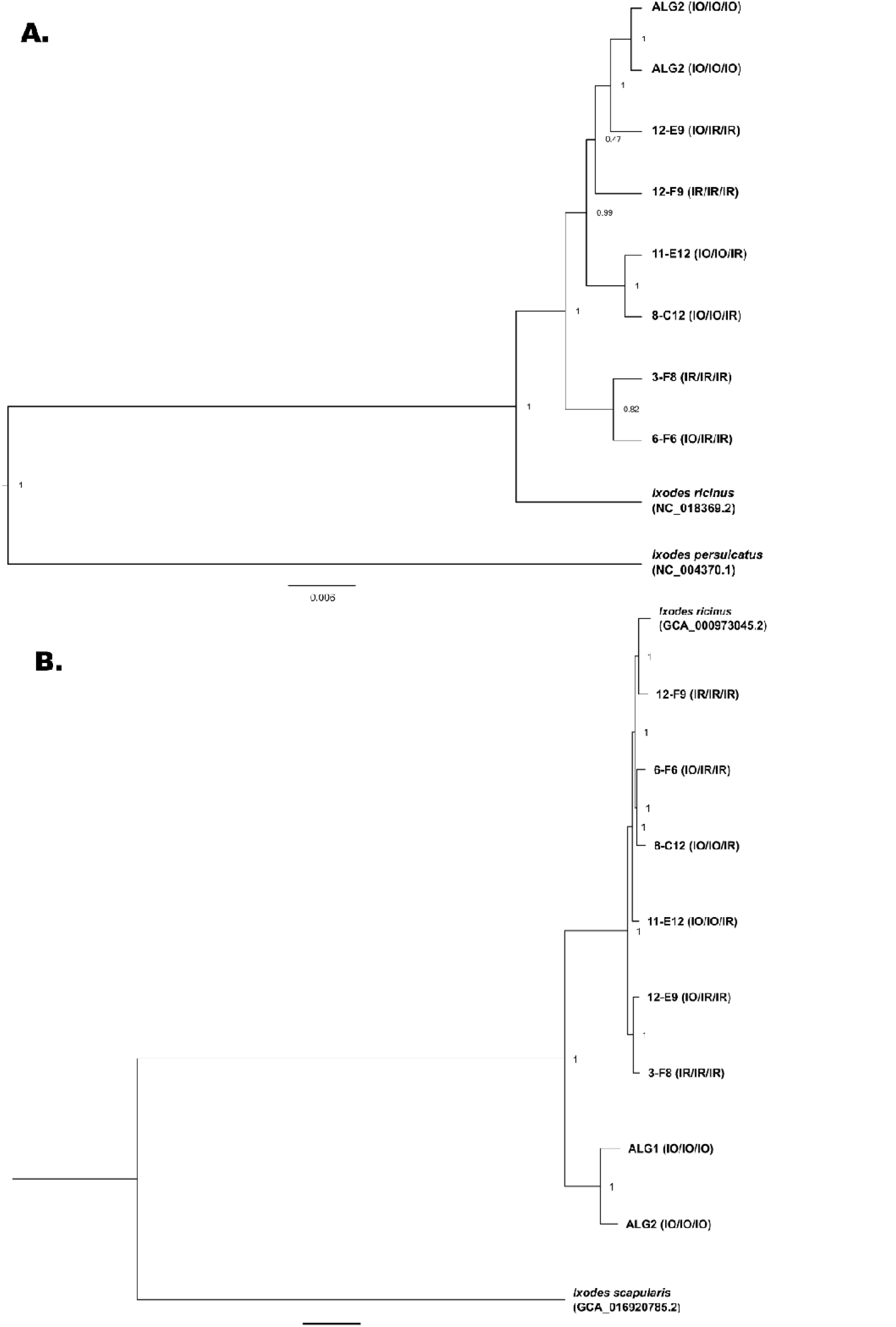
Phylogenetic reconstruction of all sequenced tick specimens based on the complete mitochondrial genome (A) or 194 unphased, nuclear sequences each of 1kb extracted from across the entire genome (B). A) The phylogeny was estimated in BEAST v2.6.6 (Bouckaert et al., 2019) with the following parameters: gamma site model with four categories, log-normal relaxed clock (Drummond et al., 2006), GTR substitution model (Tavaré, 1986). B) Phylogeny reconstruction was performed in MrBayes v3.2.6 as a partitioned analysis (Huelsenbeck and Ronquist, 2001; Ronquist et al., 2012). For this, each gene was allowed to evolve independently by estimating a GTR (Tavaré, 1986) substitution model with inverse gamma distributed rate variation and ploidy set to diploid per individual gene. For both trees, three independent runs were launched for 50 million (A) or 100 million (B) generations. Information contained in paraentheses referes to the three criteria used to choose samples for whole genome sequencing corresponding to morphological identification, 16S sequences, and TROSPA sequences. IO refers to Ixodes inopinatus and IR to Ixodes ricins.

### 3.3. Estimation of demographic history based on nuclear sequences

All tick specimens showed similar heterozygosity based on reconstructed nuclear sequences, but Algerian *I. inopinatus* samples showed slightly higher heterozygosity (Figure 3). Additionally, k-means clustering on the individually phased sequences (see Supplementary Methods) showed that the proportion of sequences where individual haplotypes belonged to different clusters was not higher in any of our sequenced samples (Figure S2). In this analysis, all reference sequence haplotypes for *I. ricinus* and *I. scapularis* always belonged to the same k-means cluster (Figure S2).

**Figure 3.**
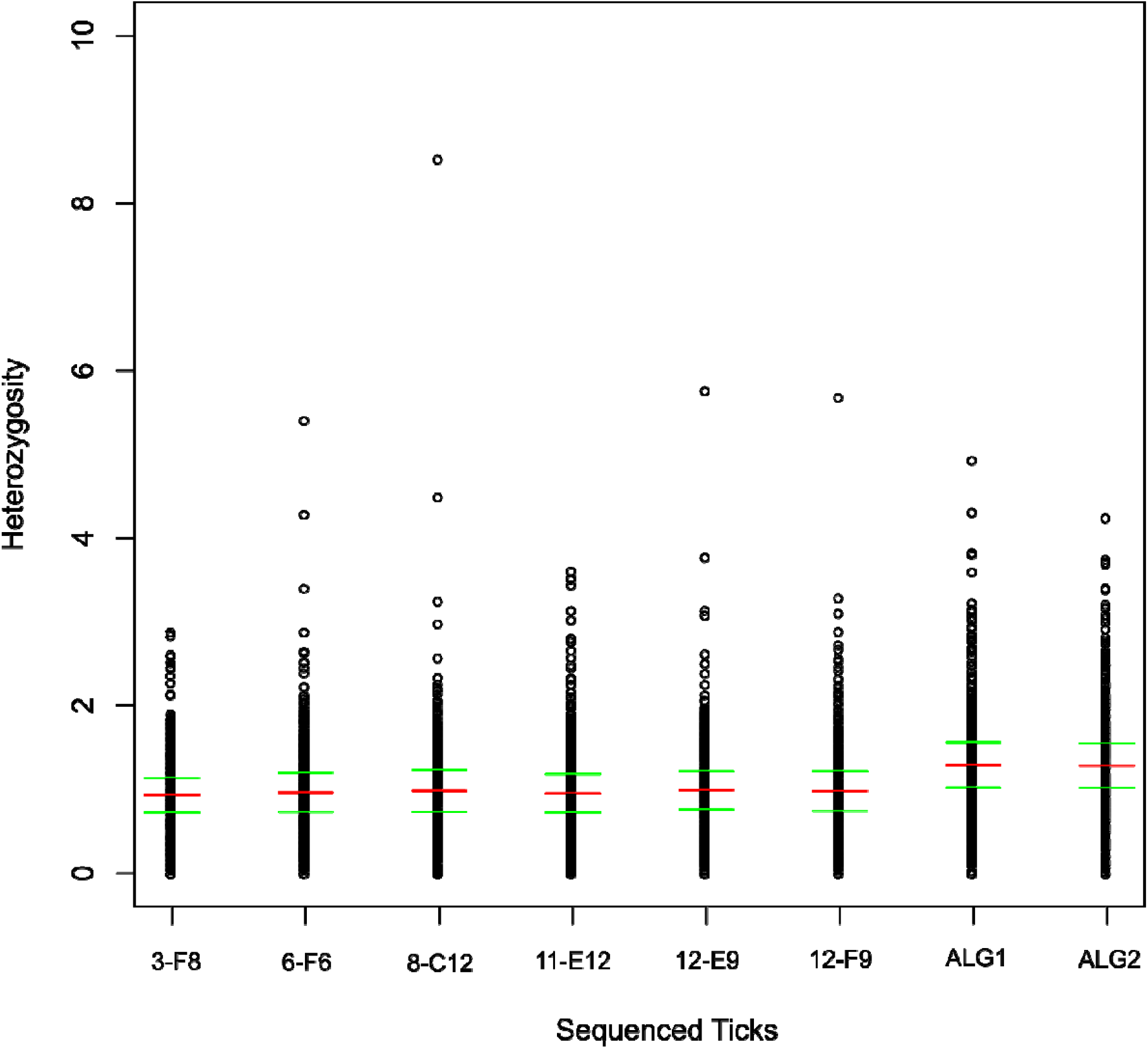
Estimated heterozygosity of all sequenced tick samples based on 435 loci distributed across the genome. Heterozygosity was estimated in R v.3.5.2 (R Core Team, 2019) using the package GENHET (Coulon, 2010).

Demographic modeling inferred that the German *I. ricinus* (Pop 1) had a larger population size than the assumed German *I. inopinatus* (Pop 2) (*N*_*1*_*/N*_*2*_=19.13, CI: [18.82, 29.05], with the population-mutation rate per locus for population 2 estimated as θ_2_=2.58, CI: [1.34, 2.64]) (Figure 4). The Algerian *I. inopinatus* (Pop 3) also displayed a larger population size in comparison to the assumed German *I. inopinatus* population 2, but with a lower magnitude when compared to the German *I. ricinus* population (*N*_*3*_*/N*_*2=*_7.60, CI: [6.45, 10.13]) (Figure 4). The German populations were estimated to have diverged very recently in time (*t*_*12*_=1.03, CI: [1.03, 1.47]) (Figure 4) with the divergence between Algerian *I. inopinatus* and the assumed German *I. inopinatus* being more ancient (*t*_*32*_=8.14, CI: [7.56, 10.89]) (Figure 4). Models showed high migration between German populations in both directions: namely, German *I. ricinus* into the assumed German *I. inopinatus* (*M*_*12*_=9.77, CI: [9.77, 12.94]) and assumed German *I. inopinatus* into the German *I. ricinus* (*M*_*21*_=2.58, CI: [0.75, 4.71]) (Figure 4). Note that these estimated rates would imply, for example, that if two alleles of a gene locus are sampled from population 2 and their ancestral lineages are followed back in time, the rate at which one of them traces back its ancestry to population 1 is 9.77 times as high as the rate at which they coalesce in population 2. In comparison, very little migration was observed between the assumed German *I. inopinatus* (Pop 2) and the Algerian *I. inopinatus* (Pop 3) in either direction (*M*_*32*_=0.33, CI: [0.23, 0.38]; *M*_*23*_=0.17, CI: [0.08, 0.17]) (Figure 4). The recombination rate was estimated as ρ=3.23 (CI: [1.09, 3.77]) and the transition– transversion ratio as 2.7 (CI: [2.43, 2.85]).

**Figure 4.**
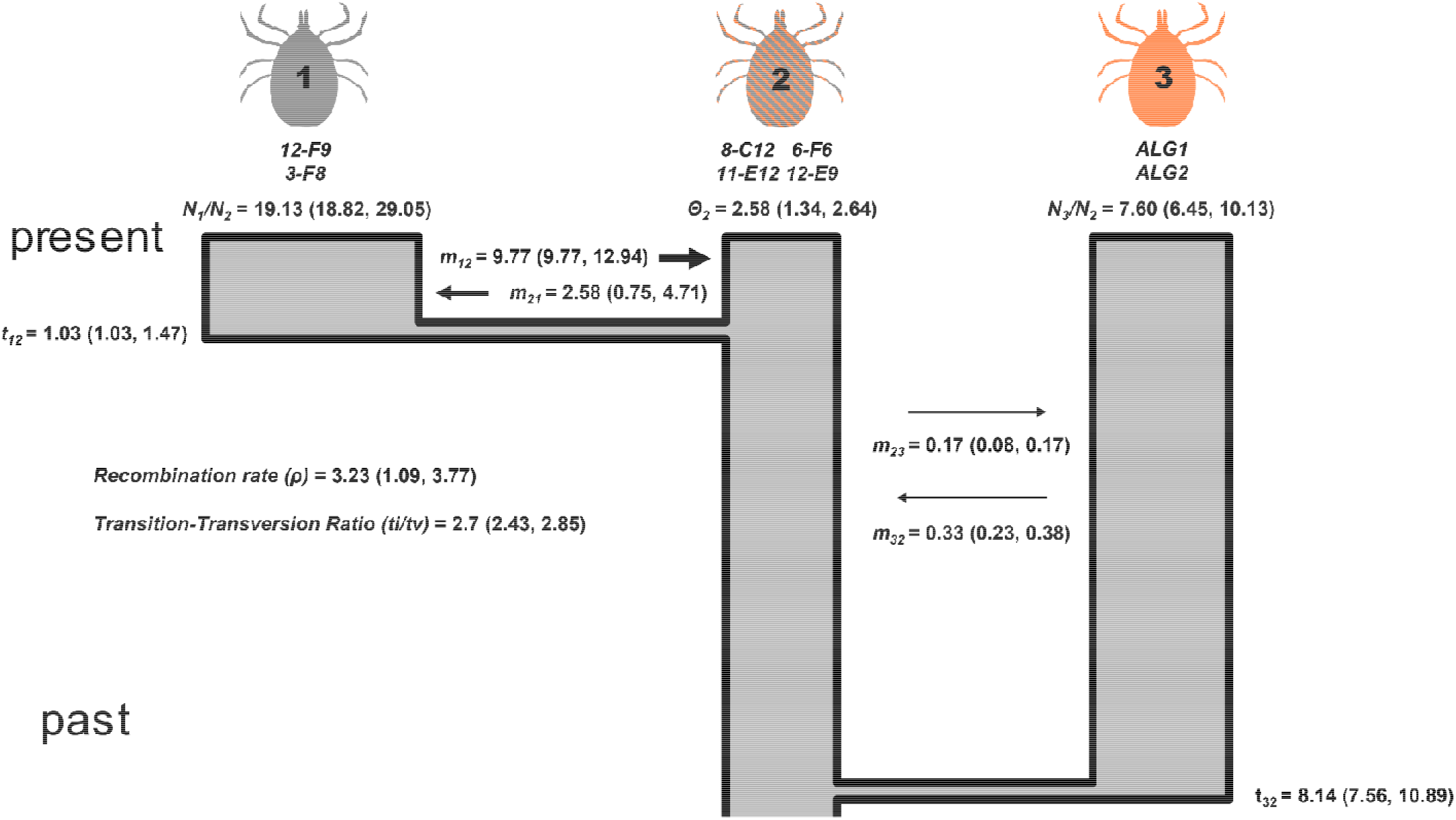
Demographic parameters estimated using Jaatha 3.2.3 (Mathew et al., 2013; Naduvilezhath et al., 2011). The model assumed three populations which all share a common ancestor: Population 1, German I. ricinus samples (12-F9, 3-F8); Population 2, German I. inopinatus samples (8-C12, 11-E12, 6-F6, 12-E9); and Population 3, Algerian I. inopinatus samples (ALG1, ALG2). We used the R package coala, version 0.7.1, to specify and simulate the demographic model in R (Staab and Metzler, 2016; https://CRAN.R-project.org/package=coala), combined with the scrm package (version 1.7.4) the seq-gen program, version 1.3.4 to generate sequences evolving along the simulated genealogies according to the HKY sequence-evolution model (Rambaut and Grass, 1997; Staab et al., 2015). Parametric bootstrap values are included in parentheses calculated from 50 repetitions (Efron and Tibshirani, 1993; Mathew et al., 2013) applying the reverse-percentile intervals method as implemented in the R command boot.ci with option type=‘basic’ (boot package version 1.3-28.1).

## 4 Discussion

Many tick species act as vectors for various human and animal pathogens (Guglielmone et al., 2014; Hillyard, 1996) but even so, unambiguous identification and unclear taxonomy hamper our ability to unravel the evolutionary history of these important arthropods. This is true in the case of two closely related species *I. ricinus* and *I. inopinatus*, which are found in sympatry over their European and North African ranges (Chitimia-Dobler et al., 2018; Estrada-Peña et al., 2014; Rubel et al., 2021; Younsi et al., 2020). During a previous study, 37 larva and nymphs were morphologically identified as *I. inopinatus* (Rollins et al., 2021), which we aimed to confirm using molecular data in this study. The results presented here display that the samples found in Germany, although morphologically similar to *I. inopinatus*, do not represent this species.

*Ixodes inopinatus* was originally described based on morphological characteristics and sequence variability along the mitochondrial 16S rRNA gene (Estrada-Peña et al., 2014). although recent work has suggested that the 16S rRNA gene alone is not adequate to delineate *I. inopinatus* from *I. ricinus* (Plantard et al., 2022). The results presented here extend this by showing that even full mitochondrial genome sequences are not able to distinguish *I. ricinus* from *I. inopinatus* samples (Figure 2A); only when considering nuclear sequences is the difference apparent (Figure 2B) as previously suggested (Noureddine et al., 2011; Poli et al., 2020). This suggests that the divergence between these species is relatively recent as the mitochondrial genome evolves much slower in comparison to the nuclear genome (Cameron, 2014; Wang et al., 2019), although future work would be needed to determine when this split occurred historically. In this study, we opted to use 16S rRNA in tandem to a nuclear gene shown to potentially delineate *I. ricinus* and *I. inopinatus* (i.e., TROSPA) (Norte et al., 2021; Noureddine et al., 2011) which we were able to support with our analysis (Figure 1B). Therefore, future research should use nuclear markers, such as TROSPA (Noureddine et al., 2011), as a viable alternative to mitochondrial genes to validate morphological identification of *I. inopinatus*.

Analysis based on morphological criteria did indeed show that these criteria can separate *I. inopinatus* from *I. ricinus* specimens (Estrada-Peña et al., 2014). We utilized the same morphological criteria to identify Algerian *I. inopinatus* samples as the assumed German *I. inopinatus* samples, and do show that, in the case of the Algerian samples, *I. inopinatus* was properly identified. Even so, the results additionally show that samples that were morphologically identified as *I. inopinatus* in Germany were in fact genetically *I. ricinus* samples when looking at a specific nuclear marker (Figure 1B) or many nuclear sequences (Figure 2B). This would suggest that the morphological characteristics are not unambiguous in delineating the two species and other methods, such as sequencing of a nuclear gene, should be considered when attempting to identify *I. inopinatus*. The fact that the assumed German *I. inopinatus* are indeed genetically *I. ricinus* could suggest that *I. inopinatus* is not present within Germany contrary to previously reported (Chitimia-Dobler et al., 2018; Hauck et al., 2020; Knoll et al., 2021). These studies used 16S rRNA sequences to confirm *I. inopinatus* morphological identification which could have led to false identification of this tick species. Previous work solely based on molecular data (no prior morphological identification of *I. inopinatus* samples) suggested also that no *I. inopinatus* samples were found outside of North Africa based on population structuring analyses (Poli et al., 2020). Our results, in addition to this previous work, could support that *I. inopinatus* is not present in Central Europe and is restricted geographically to Northern Africa and, potentially, southern Spain. This was further supported by demographic analysis, which estimated high levels of gene flow between German samples and isolation of the Algerian samples (Figure 4). This analysis also supported that *I. inopinatus* is rarer (smaller effective population size ratio) than *I. ricinus* samples (larger effective population size ratio) (Figure 4). These results taken together imply that *I. inopinatus* samples previously identified in Germany could instead by *I. ricinus* suggesting that this tick species is not present in Central Europe. Additionally, as the type specimens (collected in Spain) were confirmed through 16S sequencing (Estrada-Peña et al., 2014), these and other samples (Chitimia-Dobler et al., 2018; Hauck et al., 2020; Knoll et al., 2021) should be sequenced for additional genes (e.g., TROSPA) to confirm their identity as *I. inopinatus*.

This, however, does not explain why ticks which are genetically *I. ricinus* appeared to be *I. inopinatus* based on morphological characteristics. A potential explanation could be that the German samples morphologically identified as *I. inopinatus* represent *I. ricinus/I. inopinatus* hybrids as *Ixodes* species are known to naturally hybridize (Bugmyrin et al., 2016; Kovalev et al., 2016, 2015). Previous work utilizing TROSPA to identify *I. inopinatus* supported their presence on migratory birds traversing into Europe (Toma et al., 2021). This could potentially allow for movement of *I. inopinatus* samples into Europe further allowing for a chance of hybridization with local *I. ricinus*. Demography results presented here do also show a small amount of gene flow between German and Algerian samples suggesting hybridization could occur (Figure 4). Even so, German samples originally identified as *I. inopinatus* do not display higher heterozygosity (Figure 3), as would be expected for hybrids. Moreover, phased haplotypes of these samples do not sort to different groups in a k-means clustering analysis more often than other sequenced samples (Figure S2). These data along with placement within the nuclear phylogeny (Figure 2B) would suggest that these samples do not support the hypothesis that *I. ricinus*/*I. inopinatus* are hybrids. Future research would need to be done including mating experiments to determine if these ticks can hybridize and if phenotypic characterization of the hybrids corresponds to wild ticks captured in Europe morphologically identified as *I. inopinatus*.

The results presented here highlight the difficulty in separating out closely related tick species with clear separation only being achieved through sequencing nuclear markers. Previous work has mentioned similarities between the clustering of *I. scapularis* in North America and the clustering of *I. ricinus* and *I. inopinatus* between Europe and North Africa (Noureddine et al., 2011). Indeed, *Ixodes scapularis* exists in two distinct lineages along the east coast of North America: Clade A (northern clade) and Clade B (southern clade) (Qiu et al., 2002; Sakamoto et al., 2014). This brings forward the question if *I. ricinus* and *I. inopinatus* could truly represent two geographically separated populations of one species. This of course would need to be tested through mating experiments and using larger nuclear datasets including samples from many sampling locations. Either way, the analysis reported here shows the power of utilizing whole genome sequencing data in unraveling open questions within the evolutionary history of ticks and provides insight into how best to tease apart closely related species reliably in future studies.

## 5 Acknowledgments

We would like to thank all past members of the Evolutionary Biology group at the LMU as well as Alexander Graf from the LMU Gene Center, all lab technicians and colleagues at the National Reference Centre for *Borrelia*, all members of the Behavioural Ecology group at LMU. Additionally, we would like to thank Dr. Chelia Houcine for help in locating suitable sampling sites in Algeria.

## 6 Author contributions

RER, NSB, LCD, and VF conceptualized the project. Samples were collected by RER, AM, ND, AL, NR, and LCD; including morphological identification which was done by RER and LCD. Lab based data collection and sequencing were organized and run by RER, AB, SK, and LCD with guidance and assistance from GM, VF, NSB. NSB, AB, and SK assembled and produced all genomic data and analyses on this data were performed by DM (demography), RER (phylogenetics, heterozygosity), and NSB (phylogenetics, heterozygosity). RER wrote the manuscript with the help of NSB, LCD, and DM. The final manuscript was read and approved by all co-authors.

## 7 Research funding

The project was funded through the German Research Foundation (DFG Grant No. BE 5791/2-1) (NSB, RER). The National Reference Centre for *Borrelia* was funded by the Robert-Koch-Institut, Berlin (VK, GM). ND was funded by the German Research Foundation (DI 1694/1-1).

## 8 Ethics statement

All ethical approval and ethics committee permissions for animals used in this study was granted by the District Government of Upper Bavaria (Regierung von Oberbayern) for Animal Care (Permit Number: ROB-55.2-2532.Vet 02-17-215) in accordance with the ASAB/ABS Guidelines for the use of animals in research.

## 9 Data availability

All molecular data (16S rRNA, TROSPA, and SRA files) are currently being uploaded to GenBank and will be made available upon publication of this manuscript.

