## Supplementary Materials for "German *Ixodes inopinatus* samples may not actually represent this tick species"

**International Journal for Parasitology**

Robert E. Rollins, Gabriele Margos, Andreas Brachmann, Stefan Krebs, Alexia Mouchet, Niels J. Dingemanse, Abdelkarim Laatamna, Nassiba Reghaissia, Volker Fingerle, Dirk Metzler, Noémie S. Becker, and Lidia Chitimia-Dobler

- **Online Supplementary Material —**
- **Supplementary Methods —**

*Determining clustering based on phased nuclear sequences*

The aligned nuclear genome sequences with outgroup (i.e., *I. scapularis*) (n=194) were individually phased using PHASE v.2.1 (Stephens et al., 2001; Stephens and Donnelly, 2003) as implemented in DnaSP v.6 (Rozas et al., 2017). For each gene, a distance matrix was calculated using the *dist.dna* command from the package “ape” (Paradis and Schliep, 2019) in R v.3.5.2 (R Core Team, 2019). A k-means clustering analysis using the *kmeans* function with k=3 from the base R “stats” package (R Core Team, 2019) was then run on each individual distance matrix and a score was given per gene if the two phased haplotypes belonged to the same (0) or different k-means clusters (1). The proportion of genes having haplotypes with the same or different cluster was then calculated for each sequenced tick individual. K was set to three in these analyses as three clusters corresponding to the three tick species (*I. inopinatus*, *I. ricinus*, *I. scapularis*) were expected *a priori*.

*Checking linkage for demographic modeling*

The summary statistics that we have used for fitting the model parameters to the data were coarsenings of the joint site-frequency spectra of all pairs of populations, resulting in 23 summary statistics for each of the three pairs of populations. To obtain summary statistics that are sensitive to aspects linkage, we checked within each of the 194 loci which pairs of polymorphisms violated the four-gamete condition for each possible phasing of the data. We calculated their fraction *x* among all pairs of polymorphisms with distance of 2 to 100 and their fraction *y* among all pairs of polymorphisms of distance 101 to 999. As additional summary statistics we then used the numbers of loci with *x*≤0.1 and *y*≤0.1, those with 0.1<*x*≤0.3 and *y*≤0.1, those with 0.3<*x* and *y*≤0.1, those with x≤0.1 and 0.1<y≤0.3, those with 0.1<x≤0.3 and 0.1<y≤0.3, those with 0.3<x and 0.1<y≤0.3, those with x≤0.1 and 0.3<y, and those with 0.3<x and 0.3<y. As a summary statistic that is sensitive for the transition–transversion ratio, we grouped the polymorphisms according to the involved nucleotide pairs and counted those for which the pair was (A, G) or (C, T) separately from the others. Further we also used the number of polymorphisms with more than two nucleotides as another statistic.

- **Supplementary Figures —**

**Table S1.** Sequencing statistics for all next generation sequencing produced.

| **Sample** | **Platform** | **Raw Reads** | **Effective (%)** | **Error (%)** | **Q30 (%)** | **GC (%)** |
| --- | --- | --- | --- | --- | --- | --- |
| **8-C12** | NovaSeq | 1115092872 | 100 | 0.03 | 92.59 | 47.2 |
| **11-E12** | NovaSeq | 1038483064 | 100 | 0.03 | 92.63 | 46.44 |
| **12-F9** | NovaSeq | 972893212 | 100 | 0.03 | 93.13 | 45.1 |
| **3-F8** | NovaSeq | 1151593050 | 100 | 0.03 | 92.41 | 46.57 |
| **6-F6** | NovaSeq | 1038528006 | 100 | 0.03 | 93.04 | 46.68 |
| **12-E9** | NovaSeq | 1016502284 | 100 | 0.03 | 93.03 | 45.51 |
| **ALG1** | MiSeq | 40938628 | - | - | 39.62 | 45 |
| **ALG1** | NovaSeq | 434068336 | 100 | 0.03 | 87.72 | 46.22 |
| **ALG2** | NovaSeq | 462582310 | 100 | 0.03 | 87.88 | 45.84 |

**Figure S1.** Schematic overview of the demographic model including estimated parameters for the three proposed populations: Population 1, German *Ixodes ricinus* (12-F9, 3-F8), German, *Ixodes inopinatus* (8-C12, 11-E12, 6-F6, 12-E9), Algerian *Ixodes inopinatus* (ALG1, ALG2).

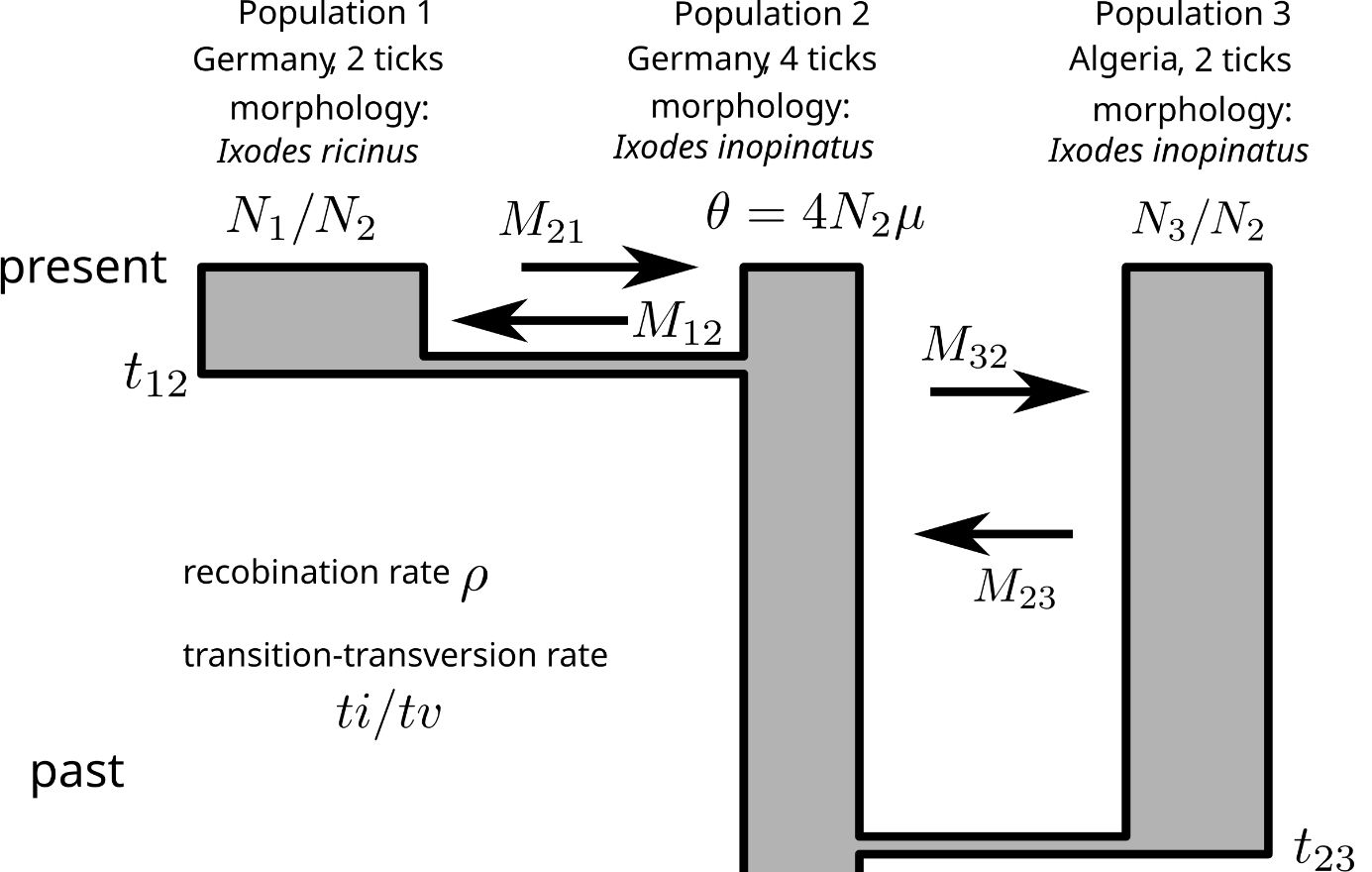

**Figure S2.** 16S rRNA alignment for all samples used in whole genome sequencing analyses including *Ixodes ricinus* and *Ixodes inopinatus* reference samples used in the first description of *I. inopinatus* (Estrada-Peña et al., 2014). Sequences were aligned using MUSCLE v3.8.425 (Edgar, 2004a, 2004b) as implemented in Aliview v1.28 (Larsson, 2014). Alignment was visualized using ESPript v. 3.0 (Robert and Gouet, 2014). Alignment continued on following page.

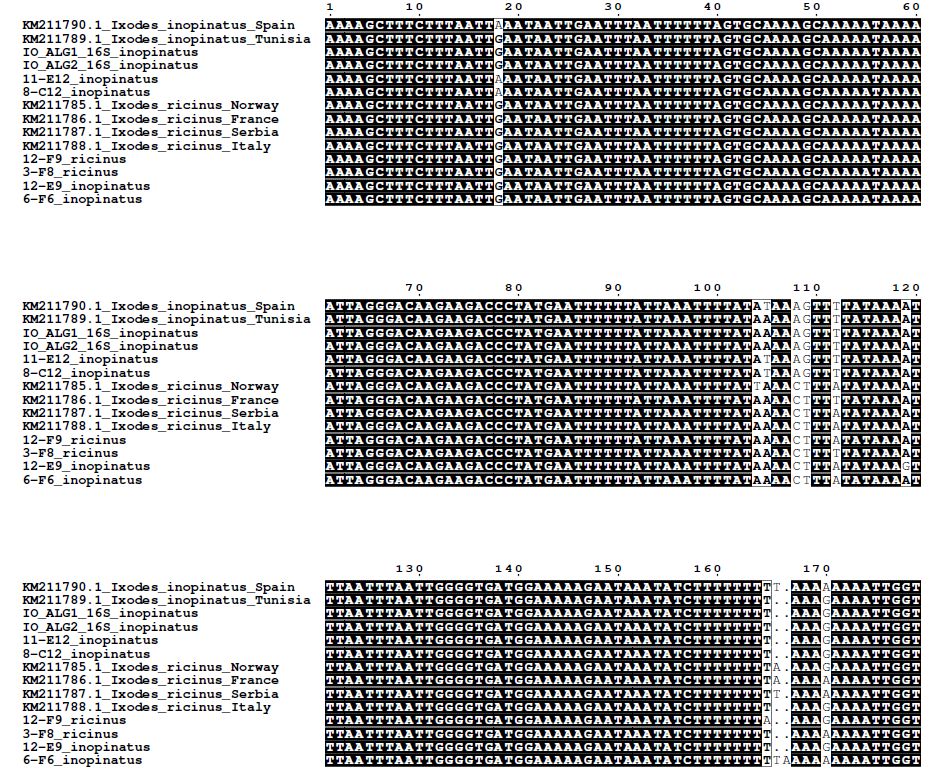

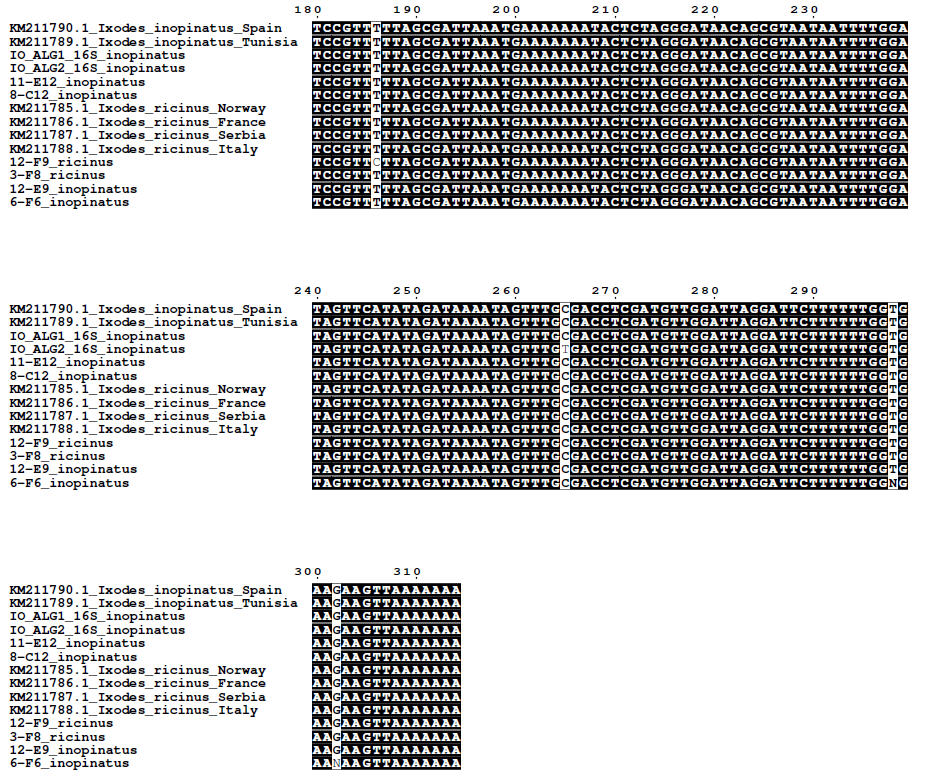
**Figure S2.** (cont.)

**Figure S3.** Proportion of phased haplotypes belonging to the same or different k-means cluster as estimated from a distance matrix calculated per individual phased nuclear gene (n=194) using the *dist.dna* command from the package “ape” (Paradis and Schliep, 2019) in R v.3.5.2 (R Core Team, 2019). K-means clustering analysis was run using the *kmeans* function with k=3 from the base R “stats” package (R Core Team, 2019). K was set to 3 as three clusters corresponding to the three tick species (*I. inopinatus*, *I. ricinus*, *I. scapularis*) were expected.

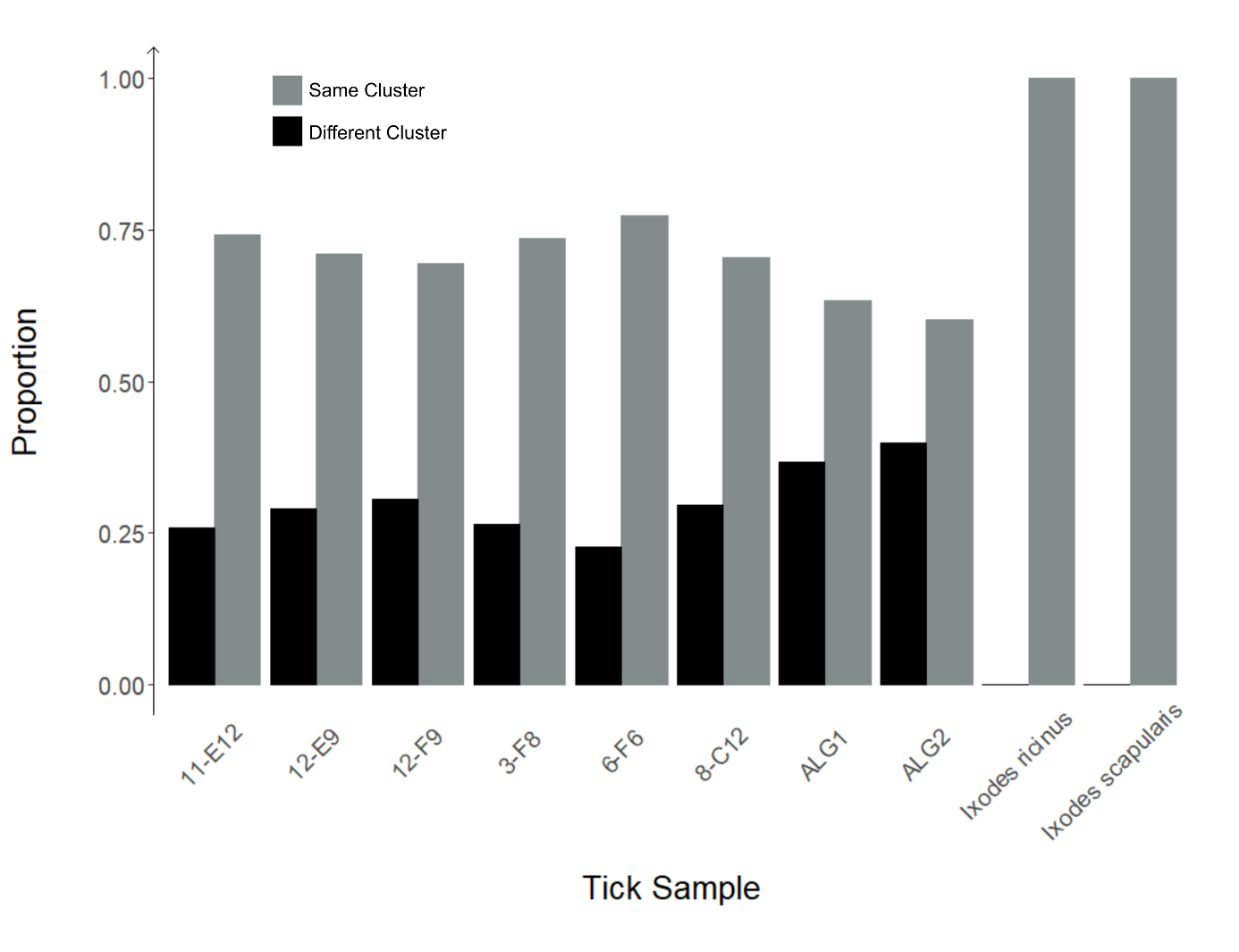
